# Fine-scale measurement of the blind spot borders

**DOI:** 10.1101/2022.07.13.499892

**Authors:** Annegret Meermeier, Markus Lappe, Katharina Rifai, Siegfried Wahl, Michele Rucci

## Abstract

The blind spot is both a necessity and a nuisance for seeing. It is the portion of the visual field projecting to where the optic nerve crosses the retina, a region devoid of photoreceptors and hence visual input. The precise way in which vision transitions into blindness at the blind spot border is to date unknown. A chief challenge to map this transition is the incessant movement of the eye, which unavoidably smears measurements across space. In this study, we used high-resolution eye-tracking and state-of-the-art retinal stabilization to finely map the blind spot borders. Participants reported the onset of tiny high-contrast probes that were briefly flashed at precise positions around the blind spot. This method has sufficient resolution to enable mapping of blood vessels from psychophysical measurements. Our data show that, even after accounting for eye movements, the transition zones at the edges of the blind spot are considerable. On the horizontal meridian, the regions with detection rates between 80% and 20% span approximately 25% of the overall width of the blind spot. These borders also vary considerably in size across different axes. These data show that the transition from full visibility to blindness at the blind spot border is not abrupt but occurs over a broad area.

## INTRODUCTION

The punctum caecum or blind spot is a disruption of visual input. Located approximately 15 to 20 deg in the temporal periphery, it corresponds to the region on the retina at which the optic nerve leaves the eye and no photoreceptors exist. It has a roughly oval shape given by the optic disc, the optic nerve head (Armaly, 1969). This region is often assumed to be small, but it actually spans a considerable fraction of the visual field, more than 50 times the area covered by the foveola—the high-acuity portion of the retina. The reported dimensions vary extensively between studies and participants (Rhodes, 2013), ranging from 5 to 7 deg on the horizontal meridian and from 7.5 to 10 deg on the vertical meridian (Armaly, 1969; Chamlin, 1960; Dolderer et al., 2006; Rhodes, 2013; Safran et al., 1993).

Although considerable individual differences are to be expected given the large variability in retinal anatomy, a notable factor that may inflate deviations across measurements is lack of a rigorous method for estimating the blind spot edges. Measuring the transition from full visibility into blindness is complicated by several factors. A major challenge is provided by eye movements, as humans are incapable of maintaining steady gaze (Rolfs, 2009; Kowler, 2011; Rucci & Poletti, 2015). Two types of eye movements are present during fixation: small saccades, known as microsaccades, occur every few hundreds of milliseconds, normally moving both eyes (Krauskopf et al., 1960; Fang et al., 2018) toward nearby objects of interest (Ko et al., 2010; Poletti et al, 2020). In the intervals between saccades and microsaccades, the eyes wander erratically at velocities typically under 2 deg/s, a motion that serves important visual functions (Rucci & Victor, 2015; Rucci et al, 2020). Together, these movements cover an area as large as the foveola (Cherici et al., 2012).

As the eyes move, a probe at a fixed position in space will at times fall onto the functioning retina and sometimes into the blind spot. Leaving fixational eye movements unaccounted for will result in spatial smearing of the blind spot map and prevent precise mapping of visibility in the transition zones. Modern optometric devices attempt to address this problem by applying a fixation control, which checks for blinks and inaccurate fixation. However, fixational eye movements are unavoidable and, given their size, they typically go undetected by commercial optometric systems.

Furthermore, differences in stimulus size and contrast could also easily result in discrepant estimates of the blind spot. In the visual cortex, the area of the blind spot, including its border, is represented by relatively large receptive fields (Awater et al., 2005; Fiorani et al., 1992), some of which even span the entire region (Azzi et al., 2015; Fiorani et al., 1992). Use of large stimuli may, therefore, trigger perceptual filling-in mechanisms, which in turn would affect the estimate. This is problematic as filling-in can also be elicited by narrow stimuli if they extend in space, for example a sufficiently long line thinner than 4’ (Spillmann, 2006). Furthermore, one should also be careful not to underestimate the size of the blind spot, as it may happen with stimuli sufficiently large to reach the border of the blind spot even if their center is well inside the blind area. In fact, it is already established that large, high-contrast stimuli lead to a smaller estimate of the blind spot area than small, low-contrast ones (Dolderer et al., 2006). All these judgments are further complicated by the poor visual resolution of the human visual system at the eccentricity of the blind spot, which makes it hard for participants to determine whether they see the entire stimulus or just a part of it.

The measurement procedure also exerts an influence on the estimation of transition zones. Two main methods have been applied, static and dynamic, depending on whether subjects judge the visibility of stationary probes or report the appearance/disappearance of probes moving at constant speeds. In this latter procedure, since the measurement relies on the ability of the participant to report a temporal event (e.g., the probe becomes visible traveling out of the blind spot), its resolution depends on the reaction time, which is known to differ widely both across participants and, for a given individual, across trials. Modern optometric devices attempt to correct for reaction time, hereby increasing the precision of the data (Dolderer et al., 2006), but this correction cannot eliminate individual inter-trial variability. Furthermore, kinetic protocols are complicated by the different perceptual mechanisms involved in detecting the disappearance and the reappearance of the probe. For example, predictive mechanisms may prolong visibility of the probe even when it is already within the blind spot, artificially shrinking the region of blindness (Maus and Nijhawan, 2008; Maus and Whitney, 2016).

The previous considerations emphasize the need for both (a) examining in detail how the transition between visibility and blindness unfolds at the blind spot boundaries; and (b) developing reliable methods for mapping the blind spot transition regions at high resolution. In this study, we describe an approach that relies on high-resolution eye-tracking, fine localization of the line of sight, and real-time gaze-contingent control of retinal stimulation. This method enables mapping of visibility around the blind spot with arcminute precision.

## METHODS

### Participants

Twelve subjects (20-28 years old, 3 males) participated in the experiments. All participants took part in measurements of the blind spot borders on the horizontal meridian. Two of them also participated in in measurement along the vertical meridian. All participants were emmetropic, with at least 20/25 acuity in a Snellen test. Subjects gave informed written consent and were compensated for their participation. The study was approved by the institutional review board at the University of Rochester and adhered to the tenets of the declaration of Helsinki.

### Apparatus

Retinal stabilization was provided by EyeRIS (Rucci et al., 2007; Santini et al., 2007), a custom system for gaze-contingent display that operated in conjunction with a high-resolution eye-tracker. This hardware/software system guarantees precise synchronization between eye movement data and the refresh of the image on the monitor. Eye movements were measured by means of two Dual Purkinje Image (DPI) eye-trackers. These systems compare the motion of the first and fourth Purkinje Images of an infrared beam, yielding a resolution of approximately 1’ (Ko et al., 2016). Data from 7 observers were acquired by means of a Generation 6 DPI (Fourward Technologies). They were low-pass-filtered at 500 Hz, and sampled at 1 kHz. This device has an internal time delay of 0.25 ms and root mean square noise level measured by means of an artificial eye of less than 20’ (Crane and Steele, 1985). The eye movements of the other 5 participants were measured at 340 Hz by means of a recently developed digital version of this machine, a digital Dual Purkinje Image eye-tracker (dDPI; Wu et al, 2022). In both setups, participants were seated on a chair in a dimly illuminated room, while movement of the head was minimized by using a custom-fitted, dental-imprint bite bar and a head rest. Vertical and horizontal eye position data were recorded for subsequent analysis.

### Stimuli

Visibility was probed using maximum-contrast squared dots with a side length of 2’. Probes were briefly flashed (14 ms) on a ROG Swift PG278Q Gaming Monitor (27”) at a resolution of 2560 × 1440 and refresh rate of 144Hz. The monitor was positioned at distance of 75 cm from the participants’ eye in the dDPI experiments, and at 115 cm in the DPI recordings, resulting in pixel resolution of 0.9’ and 0.6’, respectively. The longer distance in the experiments with the digital dDPI was necessary to optimize eye-tracking and avoid that background light emitted from the monitor would interfer with the tracking quality. Participants viewed the stimuli monocularly with the right eye while the left eye was patched.

### Gaze localization

Accurate localization of the line of sight was achieved by means of a gaze-contingent calibration procedure described in detail in previous publications (Poletti & Rucci, 2016). In this procedure, participants used a joypad to fine-tune the parameters of a pre-estimated calibration. They sequentially fixated on a series of fixation markers and adjusted the estimated position of the line of sight, which was displayed in real-time on the monitor. The offsets inserted by the participants were iteratively incorporated in the bilinear transformation that mapped the eye-tracker output into pixel coordinates on the display. This procedure has been shown to improve the accuracy of gaze localization by one order of magnitude, therefore enabling accurate mapping of the position of the blind spot region within the visual field. Gaze-contingent calibration was also repeated at the center of the screen after every block of 10 trials, to counteract possible misalignments due to small residual head movements and other possible causes.

### Blind spot mapping

Subjects were asked to report the appearance of probes displayed at various locations around and within the blind spot. We first made sure that participants could reliably detect the probe even at the greatest eccentricities. Full visibility of the probe at all eccentricities was necessary to exclude any possible bias due to a decline in performance with increasing eccentricity. Participants were instructed to maintain stable fixation on a marker (dotted circle) displayed at a fixed location of the display (see Figure 1a).

**Figure 1.**
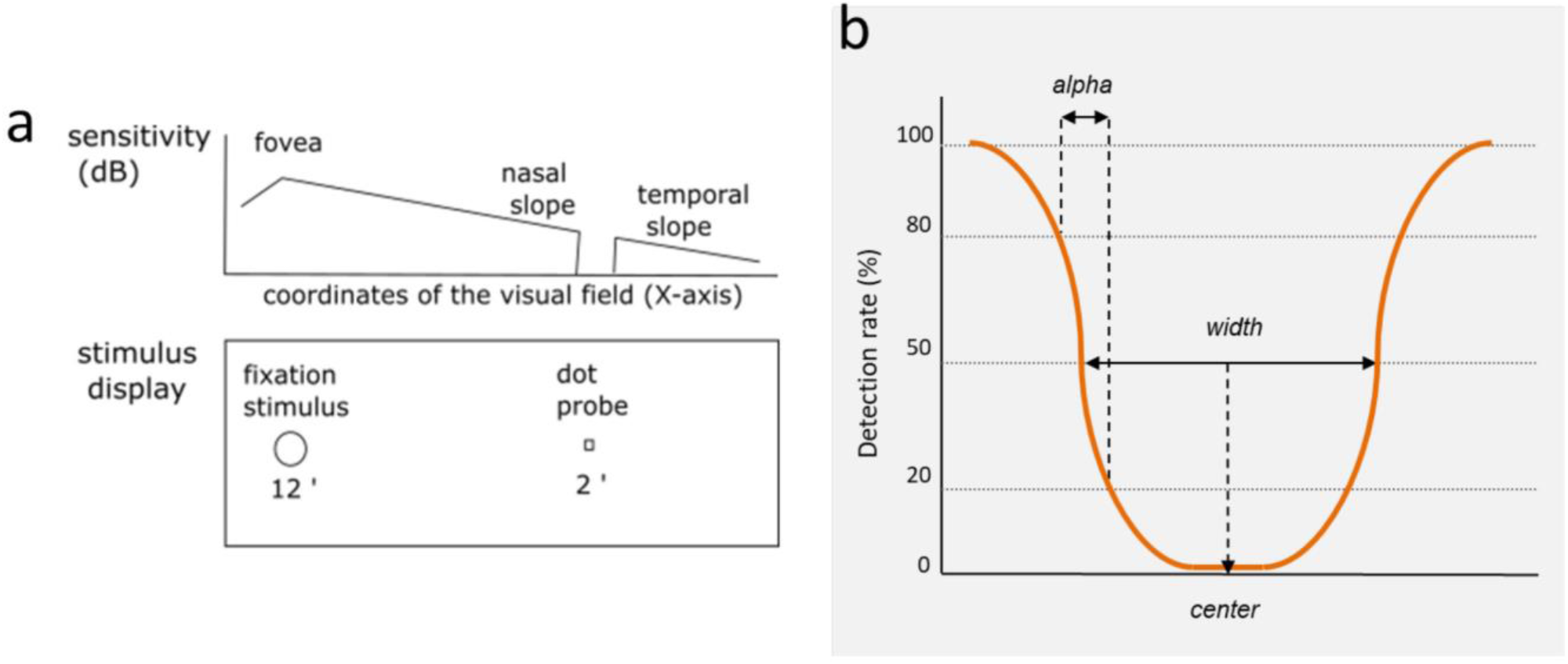
Approach and terminology used in this study. (**a**) Procedure for testing visibility around the blind spot. The upper graph represents sensitivity across the visual field. The bottom panel is a sketch of the stimulus display. Subjects maintained fixation on a 12’-diameter circle, while small high-contrast probes (2’ squares) were displayed at random locations. (**b**) Definition of the blind spot’s width, center, and border extent α. Detection rates of 20%, 50% and 80% were determined from fitted psychometric functions.

A trial started with the appearance of the fixation marker. After a variable interval of 500-1100 ms, a test probe was briefly flashed (14 ms) at the desired location. The probe was always presented under retinal stabilization, ensuring precise control of the stimulus position on the retina. The fixation marker stayed on for another 300 ms after presentation of the probe, and was then replaced by a red dot. The appearance of the red dot acted as a prompt for participants to enter their response on whether they saw the stimulus. Perceptual responses were registered via button presses on a joypad.

A coarse-to-fine procedure was used for locating the blind spot and mapping its borders. This procedure consisted of four stages, each based on the method of constant stimuli applied at a different scale. In the first stage, 12 positions equispaced by 1 deg were tested 3 times. In the second stage, 11 positions at 0.2 deg steps were repeated twice. In the third stage, 25 positions separated by just 2 pixel (1.2’ or 1.8’ depending on the setup) were presented twice. These three steps were used to provide a first approximate estimation of the location of the blind sport borders, so to narrow the area that was then examined at high resolution. This occurred in fourth stage of the procedure, when probes specifically targeted the transition zones of visibility at the margins of the blind spot. In all stages, probes were presented in random order across the possible locations. The same procedure was applied for measurements along all axes. To ensure that participants remained vigilant and complied with the task, catch trials were interspersed with normal trials. In the catch trials, the probe was flashed far outside the blind spot, at 7.5 deg eccentricity from the fovea. In the data reported in this study, participants always reported full visibility of these probes.

For two observers, to provide a quantitative example of the consequences of not considering fixational eye movements, we compared the blind spot dimensions measured in two conditions: in the presence and absence of the normal smearing of spatial measurements caused by fixational eye movements. In the latter condition, oculomotor confounds were eliminated by tracking the targeted location in real time so to present stimuli at fixed positions relative to the center of gaze.

To rigorously compare stabilized and unstabilized data, considerable number of valid trials were collected in both conditions, approximately 260 trials in each condition for one subject and ∼150 trials for the other participant.

For one observer, we systematically probed visual sensitivity in the entire space surrounding the blind spot. To this end, we used the approach described above along 60 evenly-spaced angular axes intersecting the blind spot center. Probe positions were more densely placed near the borders of the blind spot and more coarsely in the surrounding functional areas of the retina. Psychometric functions were individually estimated on each axis as to quantify the widths of the border regions along various orientations. To obtain a full two-dimensional map of visibility, performance in between the tested locations was then spatially interpolated using cubic spline functions, first along each axis and then across circular trajectories. This latter step was performed on circles with increasing radii from the blind spot center and alternating directions of interpolation (clock-wise vs. counterclock-wise) to limit artifacts in the fitting. The resulting image was then smoothed using a 6’ by 6’ median filter. Interpolation based on the original visibility data points rather than the psychometric functions obtained on each individual axis enabled better fitting of experimental data in 2D space. Overall, the map shown in Figure 6 was obtained using over 17,653 trials collected over multiple days.

For this specific subject, we also compared the results of our procedure with the anatomical data obtained by two other measurements of the blind spot: an image of the fundus of the eye and a 9 mm high-definition scan of the retina obtained via optical coherence tomography. The latter was recorded by means of a CIRRUS 5000 without additional lenses and with default settings for the fixation cross.

### Data analysis

Only periods with uninterrupted tracking and without blinks were selected for data analysis. Eye movements with speed higher than 3 deg/s and amplitude larger than 3’ were classified as possible saccadic events. If the eye velocity exceeded 2 deg/s during the 100-ms interval before and after the presentation of the probe, the trial was discarded to avoid possible errors in positioning the probe. Periods of blinks were automatically removed from data analysis. If a recalibration trial indicated a shift in accuracy in gaze localization of more than 10’, calibration was judged as having decayed substantially, and all trials acquired since the previous recalibration were discarded.

Data were analyzed using MATLAB (the Mathworks, Nattick, MA). Individual measurements were fitted with psychometric logistic functions, which were used to estimate the performance thresholds. In the following, we use the locations yielding correct detection rates of 80%, 50% and 20%. Based on these data, we estimated the width and center of the blind spot, as well as the widths of the blind spot’s borders (see Figure 1b). Specifically, the center of the horizontal blind spot was estimated as the mean between the temporal and nasal locations yielding 50% performance. The width of the blind spot was correspondingly calculated as the distance between these two points. The width of the transition zone of visibility was defined as the difference between the 20% and 80% values on the psychometric curves. This parameter is indicated by α in the following of this article. All measurements are given in angular coordinates of the visual field relative to the line of sight, so that the nasal blind spot border identifies the border closest to central fixation in the visual field. An analogous procedure was followed for processing vertical measurement data, with the axis directions oriented upward, from the lower border of the blind spot toward the upper one.

## RESULTS

The human eyes move incessantly. We are normally not aware that eye movements continually shift the projection of stimuli on the retina even during careful fixation. These movements vary widely across individuals and can cover an area as large as the entire foveola (Cherici et al, 2012). Since the spatial location of the blind spot moves together with the eye, precise mapping of the transition zones of visibility is likely affected. Yet previous studies that examined how vision declines at the blind spot edges have not considered the consequences of fixational eye movements.

To provide a quantitative example of oculomotor influences on mapping visibility, we measured the characteristics of the blind spot of two observers in the presence and absence of the spatial confound introduced by fixational instability. As shown in Figure 2a, for both subjects, there were substantial differences in the position and the width of the blind spot measured in the presence and absence of compensation for eye movements. For one participant, direct mapping of the blind spot without considering eye movements yielded a region of invisibility that was 5.11 deg wide centered at 15.22 deg eccentricity. These measurements are, however, clearly not accurate: note that failure in considering eye movements resulted in data of such low quality that the psychometric fit was actually inverted, which is physically impossible. Instead, under retinal stabilization, the borders could be measured reliably, and both psychometric functions correctly transitioned from full visibility outside the blind spot to invisibility inside. The center and width of the region measured under retinal stabilization were 15.61 deg and 5.35 deg, respectively (Figure 2b). Also in the other subject, data were poorly fitted without considering the consequences of eye movenents (Figure 2c). Without oculomotor compensation, the blind spot’s center was estimated to be 5.05 deg wide and centered at 15.45 deg. Under retinal stabilization the data were more predictable. The blind spot was now estimated to be wider and more eccentric (5.66 deg and 15.86 deg; Figure 2d).

**Figure 2.**
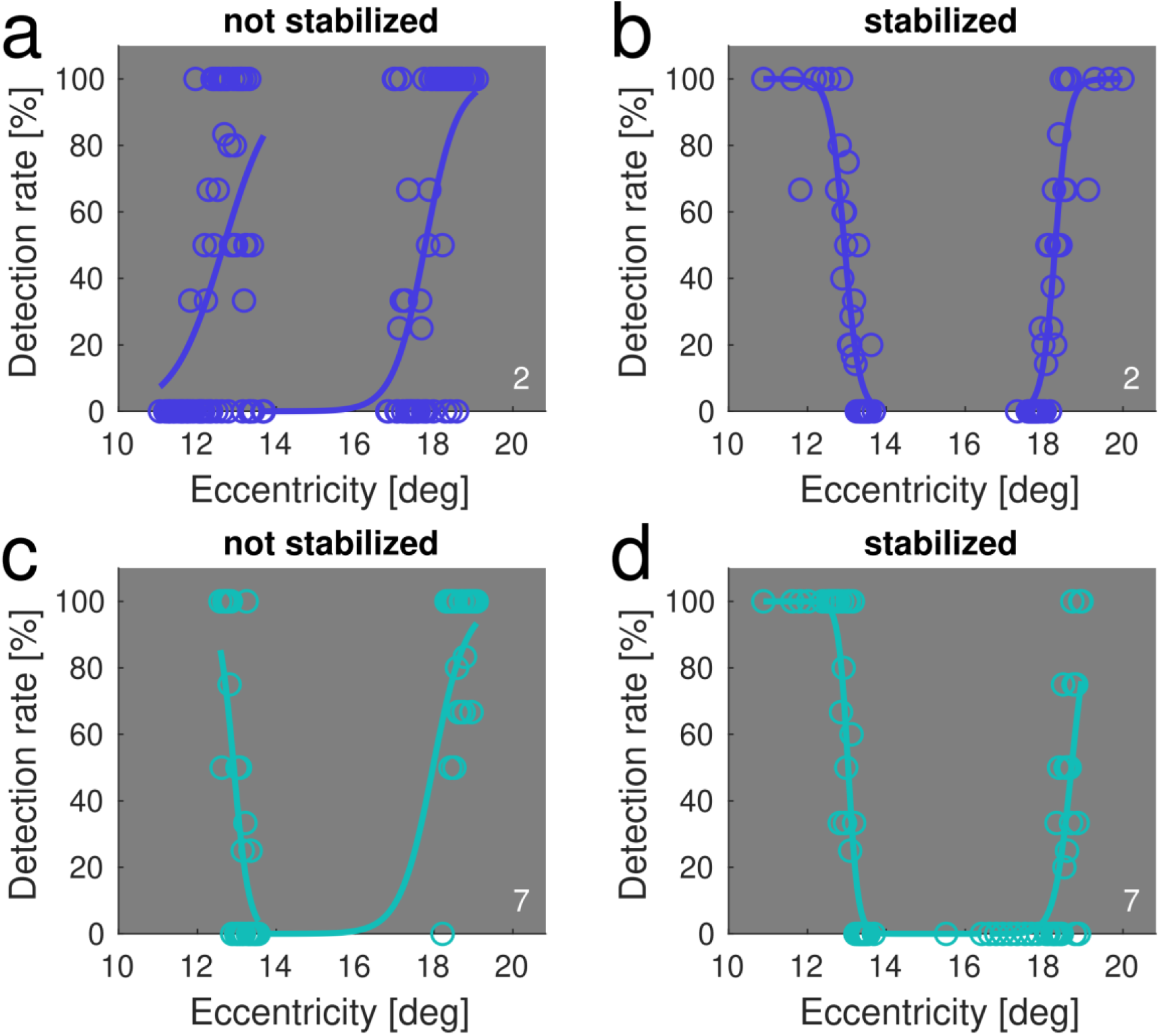
Horizontal blind spot borders measured for two participants with and without compensation for fixational eye movements. Open circles represent estimated sensitivity at various retinal locations. The solid line is the resulting psychometric fit. The two rows show data from different subjects. Measurements obtained with and without retinal stabilization are shown in the panels on the right and left columns, respectively.

The differences between measurements obtained with and without consideration of eye movements are large enough that they may affect estimation of the blind spot borders. We quantified the size of these regions via the parameter α, the distance between 20% and 80% visibility. In the first subject, the α_nasal_ and α_temporal_ measured without considering eye movements were 107’ and 32’ wide. In contrast, under retinal stabilization, these regions shrunk to 31’ and 21’, respectively. Similarly, in the second subject, α_nasal_ and α_temporal_ decreased from 67’ and 69’ without oculomotor compensation to 28’ and 37’ under retinal stabilization. Thus, on average, a 57.65% reduction was observed by properly compensating for eye movements. Stabilization also significantly improved psychometric fitting of the stabilized and unstabilized data (χ^2^(1) = 70.271, p < 0.001; binomial generalized linear model). Thus, retinal stabilization appears to be an effective method for obtaining more precise measurements and reducing uncertainty in estimating the blind spot borders.

### Horizontal borders of the human blind spot

We mapped visibility around the blind spot along the horizontal midline of the right eye for N=12 participants (Figure 3). On average, the blind spot was centered at an eccentricity of 15.53 ± 0.95 deg (mean ± standard deviation) and covered a visual angle of 5.30 ± 1.09 deg. The average widths of the transition zones of visibility at the borders of the blind spot were similar on the two sides: 0.69 ±0.50 deg on the nasal side (α_nasal_) and 0.73 ±0.38 deg on the temporal side (α_temporal_). These values that did not differ significantly (t-test for dependent samples: t(11)=-0.422, p =0.68).

**Figure 3.**
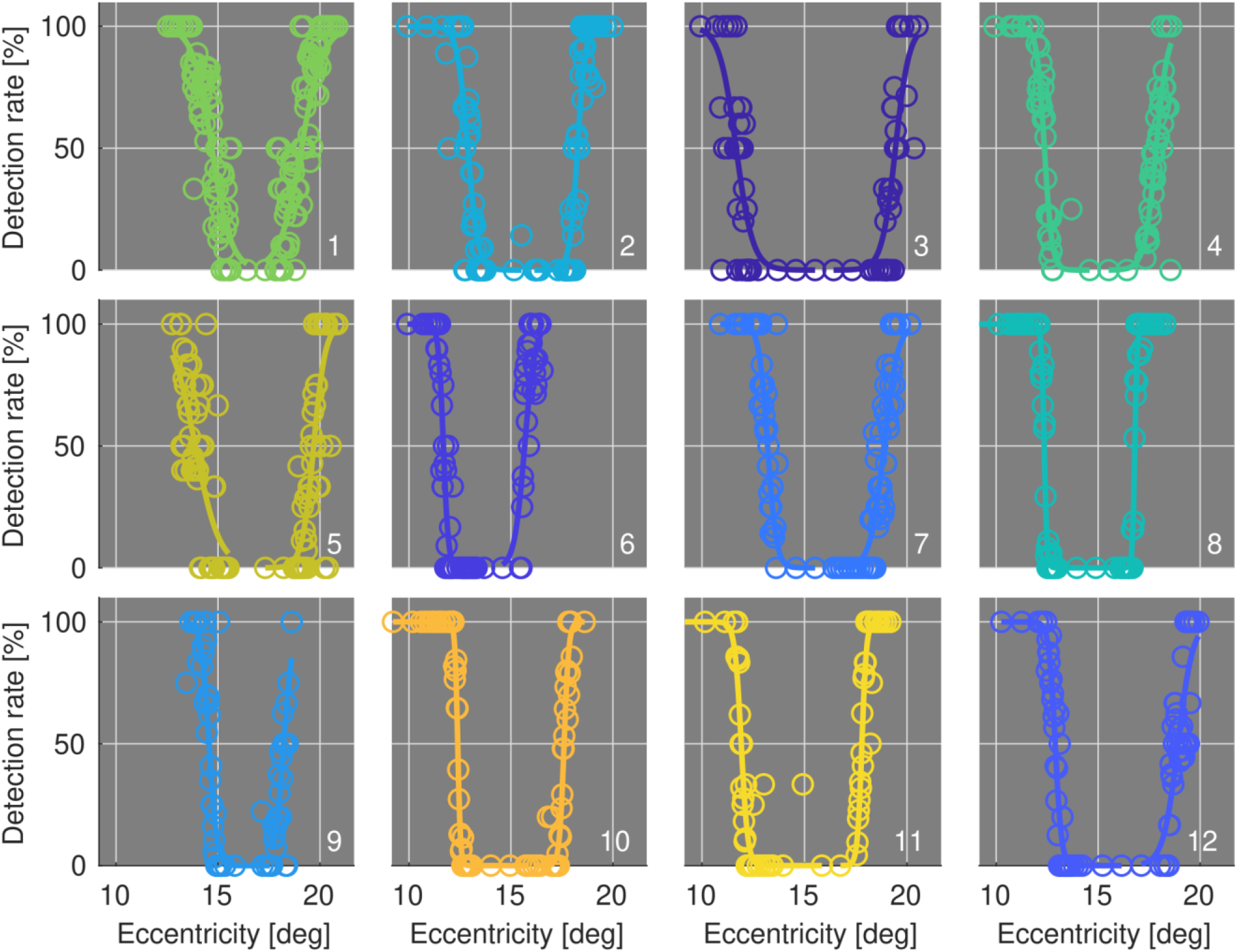
Individual measurements of the blind spot borders across the horizontal meridian. Each panel shows data from one subject. Graphic conventions are the same as in Figure 2. All data were collected while compensating for fixational eye movements via high-resolution eye-tracking and gaze-contingent display control.

Interestingly, the width of the nasal blind spot border α_nasal_ was correlated with the width of the temporal blind spot border α_temporal_ (r = 0.617, p = 0.03, Figure 4b, c), suggesting that the steepness of the blind spot borders is a feature consistent within an individual. The widths of the transition zones also exhibited a positive correlation with the eccentricity of the blind spot’s center (nasal: r =0.715, p = 0.009; temporal: r = 0.605, p = 0.04; Figure 4d). That is, the steepness of the borders decreased with increasing eccentricity of the blind spot, possibly a consequence or the declining density and increasing size of of photoreceptors.

**Figure 4.**
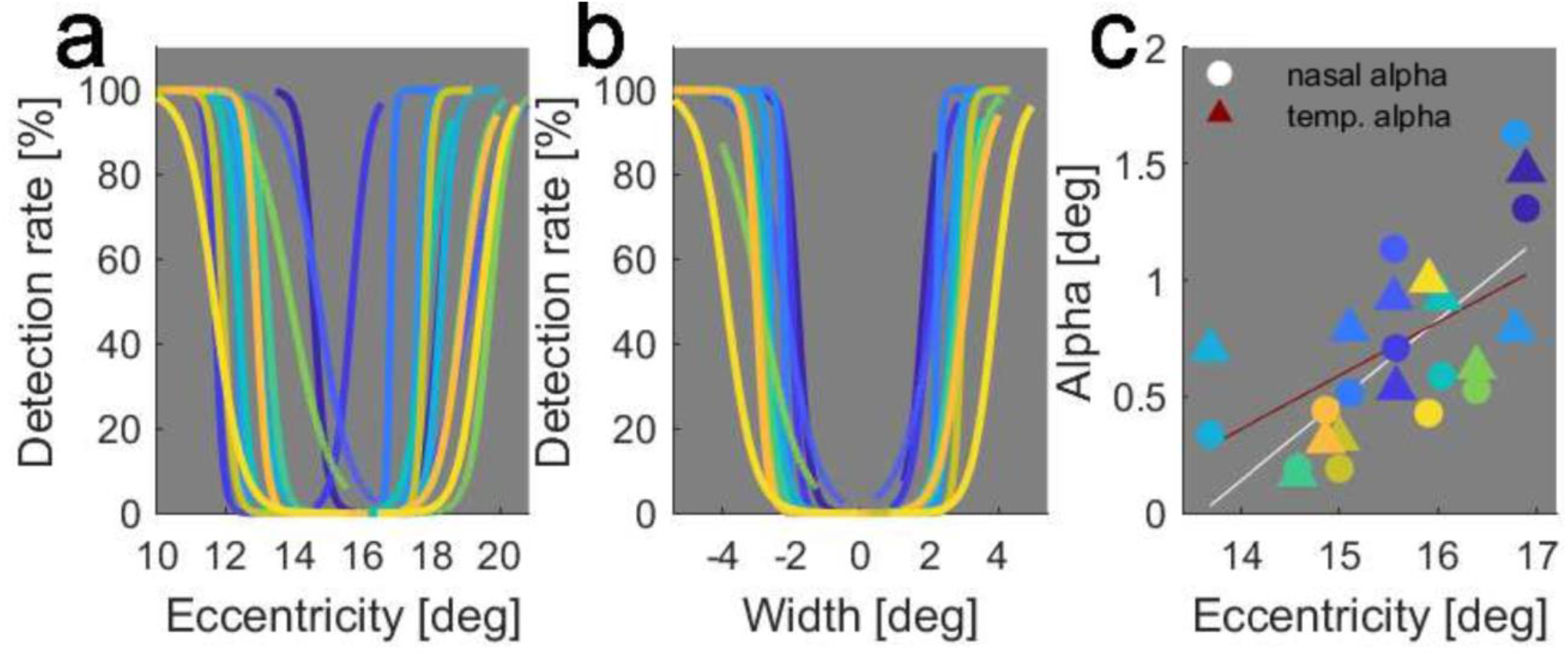
Summary of all blind spot measurements across the horizontal meridian. (**a**) Overlay of the psychometric functions of all subjects to illustrate the large individual variability in both width and position. (**b**) Same data as in panel **a**, after realigning data relative to the center position of the blind spot. Notes the similar steepness of nasal and temporal borders and the relation between the size of the blind spot and the steepness of its borders. (**c**) Correlation between widths of the nasal and temporal borders and center eccentricity.

### Vertical border of the human blind spot

In two participants, we applied our method to also measure the vertical dimensions of the blind spot (Figure 5). To this end, we first mapped visibility across the horizontal meridian and estimated the center of the blind spot on this axis. We then applied the same procedure along the vertical axis intersecting the previously estimated horizontal center. Overall, we collected 908 trials for one participant and 401 trials for the other. In the first subject, this analysis revealed that the entire blind spot was positioned quite low in the visual field and extended far below the horizontal meridian. It covered more than 11 deg in visual angle, ranging from 1.53 deg above the horizontal midline to 9.64 deg in the lower hemifield. In the other participant, the blind spot was located higher and covered a smaller extent, slightly more than 9 deg, from 2.73 deg to -6.34 deg.

**Figure 5.**
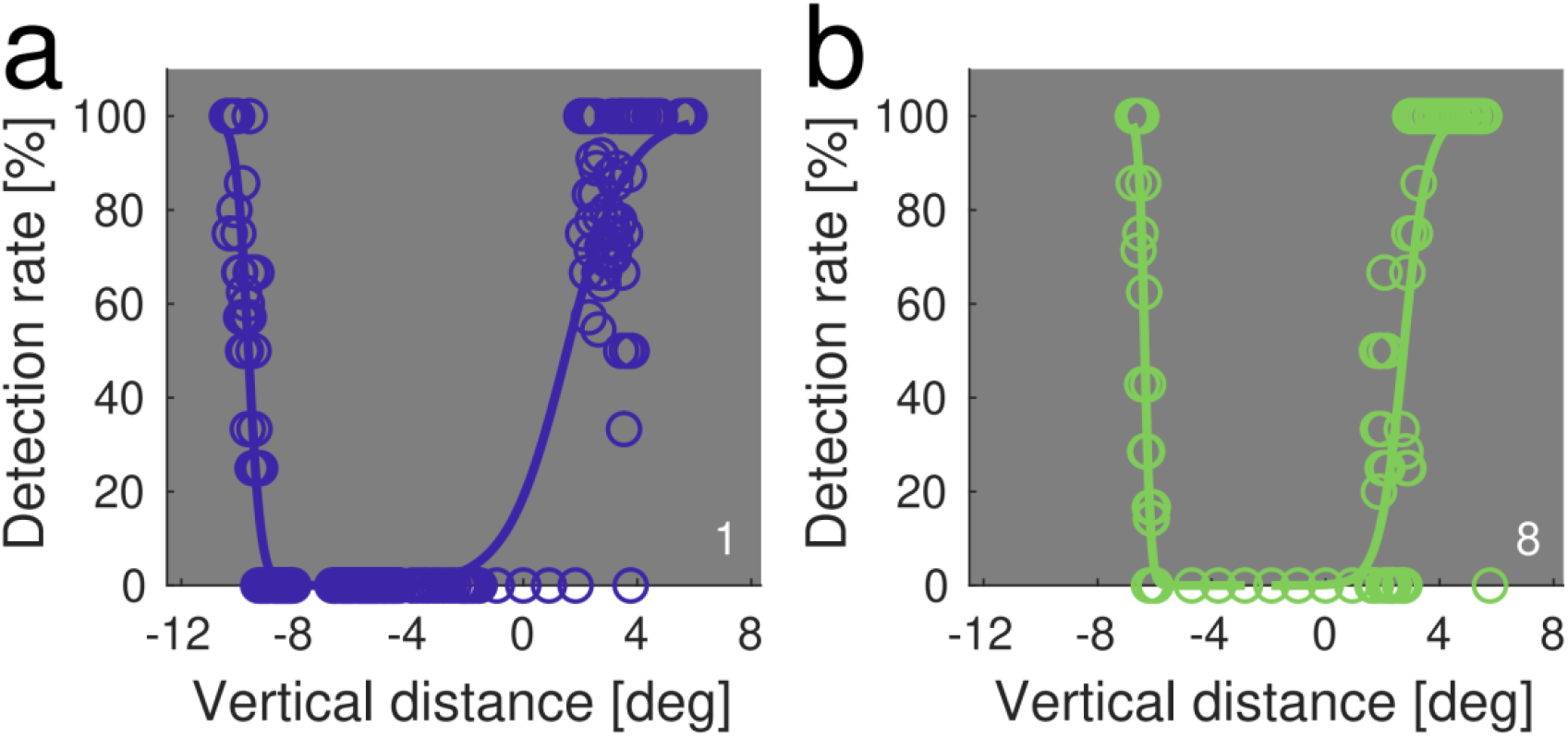
Vertical borders of the blind spot in two participants. Measurements were obtained on the vertical axis intersecting the blind spot center on the horizontal meridian, which was previously estimated. Data are plotted as a function of distance from this point.

Striking differences between the widths at the upper and lower borders were present in both subjects, with visibility transition zones much larger in the upper border. In the first subject, the transition zone only covered 36’ at the bottom but extended to almost 3 deg at the top (175’). In the other subject, the measured widths of the transition zones were 17’ at the lower border and 70’ at the upper one. It is likely that these differences were caused from angioscotomas overlapping with the vertical border of the blind spot. Major vessels typically exit the human blind spot passing from this point. This idea is supported by the 2D map of the blind spot acquired, using our method, for the subject of Figure 5b (see Figure 6).

**Figure 6.**
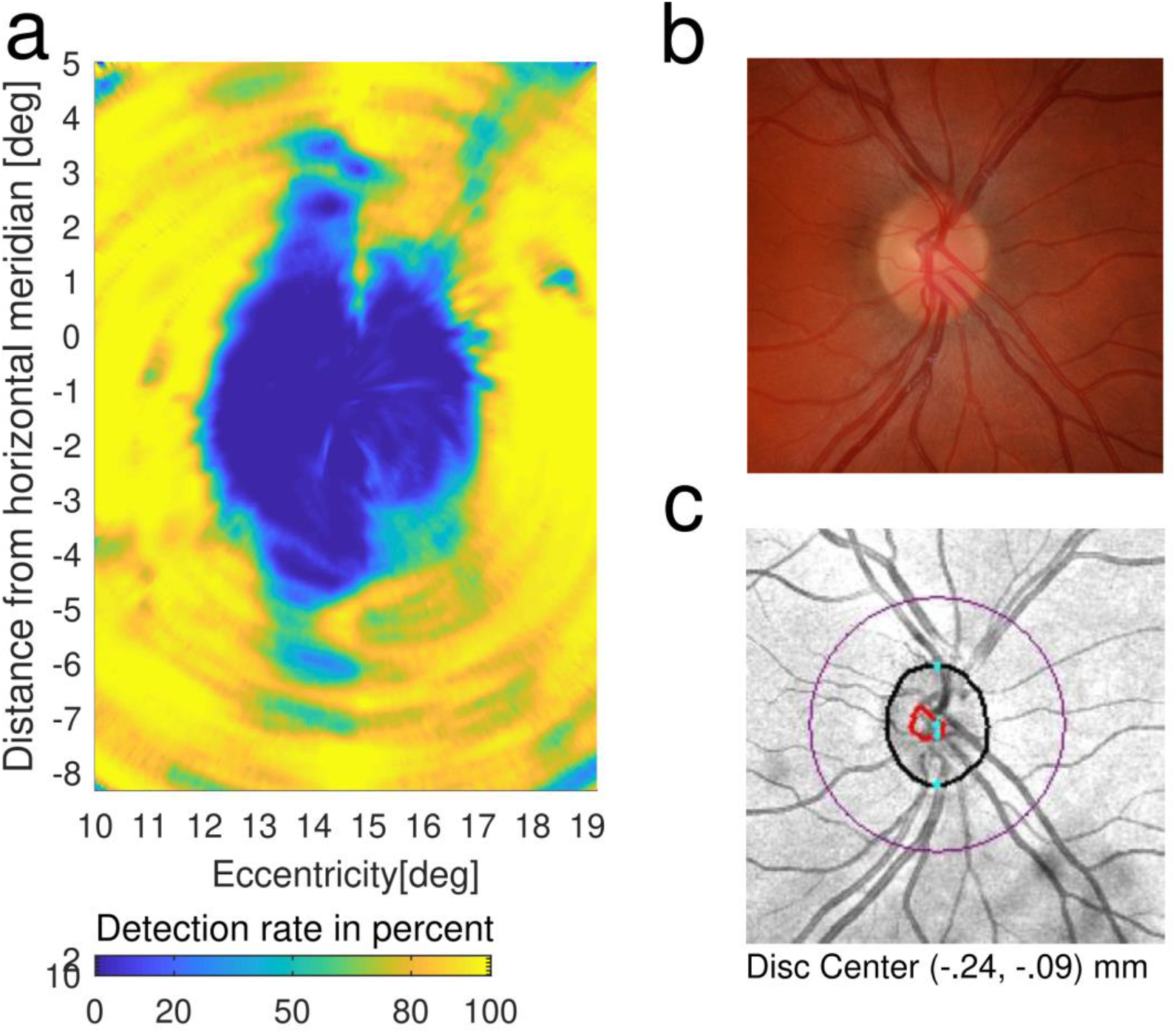
Comparison between psychophysical measurements and imaging of the blind spot. (**a**) Two-dimensional map of visibility around the blind spot for one participant. The map covers the entire blind spot region in the right eye as well as the adjacent surrounding portion of visual space. (**b**) Fundus image from the same eye showing the optic disc (light colored oval) and major vessels. Note the correspondence between the locations of visible vessels and areas of impaired visibility, suggesting that these are indeed angioscotomas. (**c**) Image of the blind spot as produced by the Cirrus 5000 HD-OCT. Dimensions are in millimeters.

### 2D map of the human blind spot

To further examine visibility around the blind spot, for one participant, we used our approach to test sensitivity across multiple axes at various orientations. This enabled derivation of a two-dimensional map of visibility with estimation of transition zones surrounding the entire blind spot. To this end, we systematically probe visibility along 60 evenly spaced angular axes. Data were then interpolated in 2D space to estimate visibility in between the tested locations.

The result in Figure 6 shows that the width of the transition zones of visibility varied substantially around the blind spot. The median width was 20’, with 25th and 75th percentiles of 14’ and 38’, respectively. This represents an average variation around the median of 61%. The width of the median nasal rim was 16’ (quartiles: 13.4’, 29.9’), whereas the width of the median temporal rim was 27’ (quartiles: 14.8’ 45.4’) a difference that was not statistically significant (z=-0.966, p =0.33; Wilcoxon sign rank test). On average, the superior edge of the blind spot extended for 89’, whereas the inferior edge was 34’ wide. On the diagonal axes, the measured widths were 30’ in the 45 deg direction, 17’ in the 135 deg direction, 15’ at 225 deg and 31’ at 315 deg. To confirm that these large fluctuations in the widths of the transition zones were partly caused by angioscotomas, for this participant we collected images of the eye fundus and compared them to our psychophysical measurements (Figure 6b). Although the two measurements differ in their reference frames (visual angles for the psychophysical measurements, retinal coordinates for the fundus image) and small rotational misalignments may be present, a strong similarity between the participant’s functional measurements and the underlying anatomical structure is clearly visible. Note that the directions which yielded broader transition zones as measured psychophysically correspond with those in which vessels are present in the fundus image. For example, the vertical borders of the blind spot closely match with the positions of major vessels leaving the optic nerve head, which are likely to be responsible for large angioscotomas.

To gain further information on the correspondence between psychophysical measurements and retinal anatomy, for this subject we also acquired a high-resolution tomographic scan of the retina using a Cirrus HD-OCT. The 9 mm horizontal image in Figure 7a shows not only the fovea but also the nasal border of the blind spot. For these images a conversion of pixels to optical angles allows registration of the functional data with the underlying substrate. The entire 9 mm image spanned a visual angle that ranged from -15.705 deg to +15.705 deg. This resulted in a factor of 0.0335 deg/pix, which we then used to overlay psychophysical measurements on the image.

**Figure 7.**
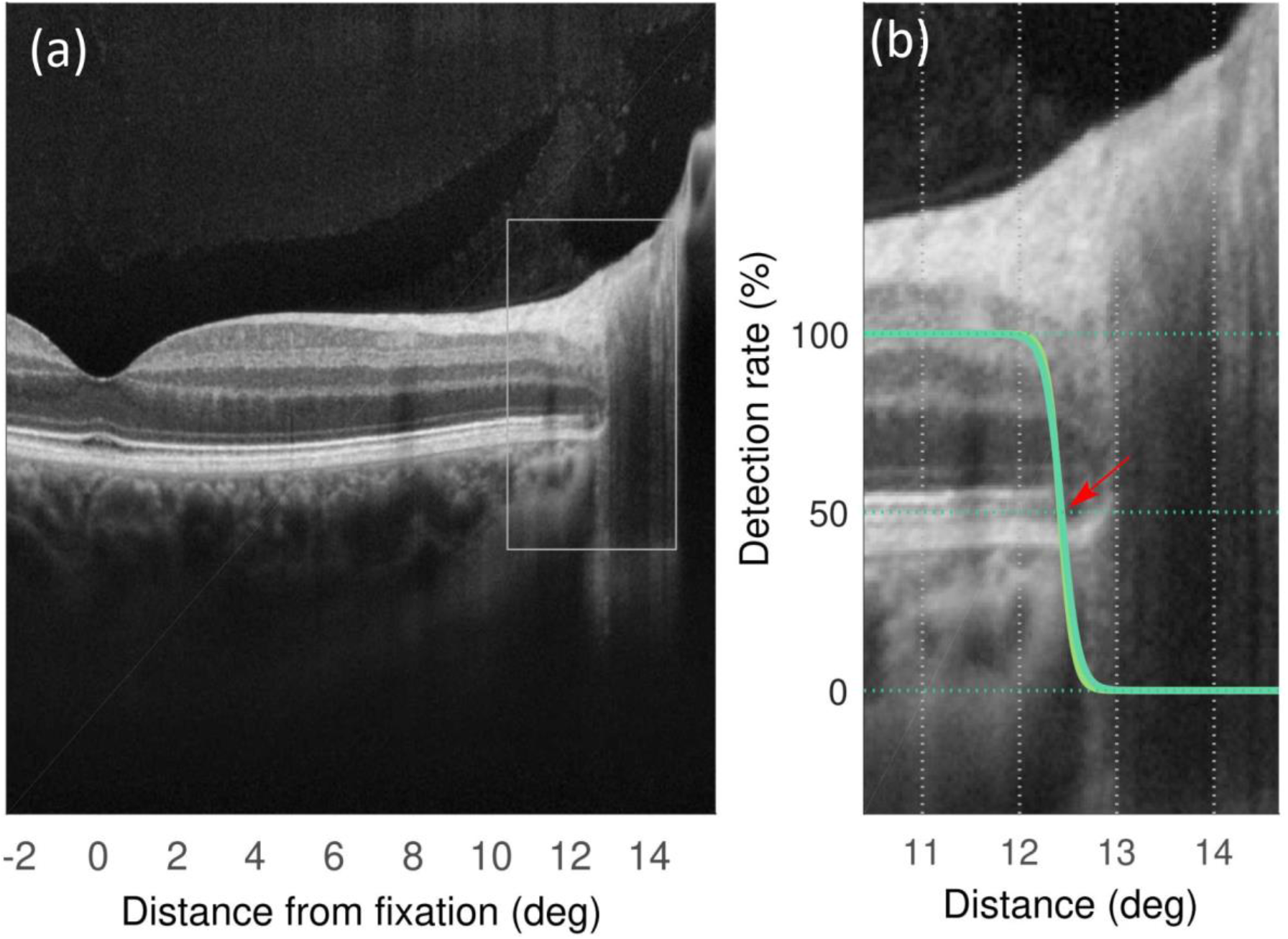
Comparison between psychophysical measurements and tomographic imaging of the retina. (**a**) 9-mm scan of the right eye of one participant (same subject as in Figure 6) using the Cirrus 5000 HD-OCT. Only the nasal portion of the scan is shown here. (**b**) Optical magnification of the retinal region at the edge of the blind spot. Superimposed is the psychometric estimation of visibility obtained using our method. The red arrow indicates the position at which the photoreceptor layer breaks off.

The image shown in Figure 7a is the retinal tomographic section corresponding to the meridian axis that was probed psychophysically. Figure 7b shows an enlargement of the specific region in which visibility was tested (gray rectangle in Figure 7a). Overlaid in green is the participant’s detection performance for stimuli around the blind spot border. As shown by these data, the location at which visibility—as tested with our paradigm—drops coincides with the anatomical locus at which the different layers of the retina, and especially the photoreceptor layers (see Figure 7 red arrow), become discontinued. These results further confirm that our approach is highly precise in estimating the location of the blind spot. Although the resolution of these data does not allow assessment of whether photoreceptors progressively decline in the transition zone, a decline in visibility appears to initiate at a location in which the photoreceptors outer segments are already disrupted. Thus, the alignment obtained in this analysis is compatible with the idea that a decline in performance is mediated by increasing axonal convergence and/or lateral connections with increasing eccentricity.

## DISCUSSION

We measured visibility around the blind spot with high-resolution perimetry under retinal stabilization. This approach compensates for the influences of the incessant eye movements that humans perform while attempting to maintain steady fixation. Since these movements shift gaze by amounts larger than the widths of the transition zones at the blind spot borders, retinal stabilization removes a critical source of measurement noise. We show that even after eliminating this oculomotor confound, the transition from visibility to non-visibility takes place over a considerable spatial extent. With the temporal and nasal borders of the blind spot each occupying about 0.7 deg of visual angle, their combined width is about 25% of the total horizontal dimension of the blind spot (about 5 deg).

The approach described in this study allows for measurement of sensitivity in the border with unprecedented accuracy. This was the result of combining several precautions. First, our method relied on probes that were highly localized in both space and time and yet reliably detected—at ceiling performance—outside the blind spot region. Probes extended for only 2’ and 14 ms and were presented at random locations to avoid possible influences of prediction mechanisms. Second, eye movements were compensated for with high precision. The resolution of the DPI eye-tracker is approximately 1 arcminute (Ko et al, 2016) and the online stabilization protocol was executed at 144 Hz. Since we discarded all trials with microsaccades and all trials with eye speed above a low threshold (120 min/s), the remaining spatial uncertainty due to eye movement was less than 1 arcminute (about 0.84’; median error 0.35’). Third, absolute measurements of eccentricity are also very accurate, as our gaze-contingent calibration improves localization of the origin of the coordinate system (the center of gaze) by approximately one order of magnitude over standard methods (Poletti et al, 2016). Thus, the transition zones from visibility to blindness that we observe at the blind spot border are unlikely to result from the residual smearing of sharp edges. They appear instead to reflect a genuine progressive decrement in visibility.

As expected from retinal anatomy, there is large variability in size and location of the blind spot across participants. Interestingly, the widths of the temporal and nasal borders both correlate with the eccentricity of the blind spot center. This indicates that, while the precise anatomical characteristics of the blind spot differ across participants, the relations among these properties remain consistent. This correlation of the border width with location is not surprising and might be related to the decrease of photoreceptor density with eccentricity and/or the eccentricity-dependent increase in the size of receptive fields of neurons in the visual system. Preliminary evidence from OCT imaging in one participant appears consistent with the latter interpretation. These images showed that the probe position corresponding to chance level detection roughly coincided with the retinal position at which the photoreceptor layer ended. However, detailed inspection indicates that detection goes below chance level shortly after the visible layer of photoreceptor is disrupted, pointing at the possible action of lateral connections in the retina or beyond. Related to this point, one might also expect the temporal border—in visual field coordinates—to be broader because of the decrease of resolution with eccentricity, since this border is located further peripherally than the nasal one. However, this was not the case in our data, suggesting that any potential difference between the two, if present, is small.

In two participants, we also determined the vertical borders of the blind spot, the axis of longest elongation (Safran et al, 1993). These data showed vast differences in the dimensions of the transition zones at the two edges. A detailed two-dimensional mapping of the blind spot in one participant confirmed that the width of the transition zone varies substantially around the blind spot. Fundus and OCT images showed that the reason for this variation is the presence of capillaries at these locations. This confirms the precision of our method, which is capable of mapping capillaries directly from psychophysical assessments.

The extent of the transition zone that we measured is much wider than the minimal frame stimulation needed to produce filling-in. Spillmann et al. (2006) showed that a rim of only 0.05 deg width placed around the blind spot suffices to produce at least partial filling-in of color. This previous study did not measure where exactly these very thin stimuli were placed, but the results suggest that partial activation of the border region might still trigger filling-in. In fact, it is possible that the gradual decrease in visibility along the border is functionally relevant to the process of filling-in, particularly if eye movements take the stimulus inside and outside the zone of visibility. Fixational eye movements could also contribute by temporally modulating the responses of neurons within the transition zone. These modulations have been shown to strongly drive visual sensitivity outside the blind spot region (Rucci et al, 2007; Mostofi et al, 2016; Intoy & Rucci, 2020), discard redundant information in natural scenes (Kuang et al, 2012; Mostofi et al, 2020) and enhance synchronous neural responses to non-redundant features in models (Rucci & Casile, 2004; Poletti & Rucci, 2008; Casile et al 2019). These previous findings raise the hypothesis that oculomotor-induced modulations within or around the transition zones could play a role in the process of filling-in, in much the same way that absence of luminance modulations leads to filling-in of the entire visual field when stationary stimuli are viewed under retinal stabilization (Ditchburn & Ginsborg, 1952; Riggs & Ratliff, 1952; Yarbus, 1967; Poletti & Rucci, 2010).

In sum, we have presented in this study a new method for precise measurement of the border of visibility that surrounds the blind spot. This method relies on the presentation of probes at desired locations under stabilization, a technique that can be flexibly implemented by coupling gaze-contingent display control with precise eye-tracking. The approach allows definition of the blind spot’s transition zone psychophysically within a few arcminutes, a level of resolution sufficient for quantitatively investigating visual functions and the mechanisms responsible for perceptual filling-in around the blind spot. The resolution of this method may also carry benefits in clinical settings. Changes in visibility around the blind spot have been reported in various conditions, including glaucoma (Coughlan & Friedmann, 1981; Su et al. 2013). Our method enables detection of small functional changes around the optic nerve, which may go unnoticed in standard anatomical exams.

## ACKNOWLEDGEMENTS

This work received funding from the European Union Horizon 2020 research and innovation program (Marie Skodowska-Curie grant agreement No 734227) and National Institutes of Health grant EY018363.

## Notes

### Competing Interest Statement

The authors have declared no competing interest.

